# Enabling predictive modeling of molecular connections between fermented foods and human inflammation through a paired dataset of cell-based models and multi-omics approaches

**DOI:** 10.64898/2026.09.22.753573

**Authors:** Elizabeth A. McDaniel, Matthew Schertler, Chantle Edillor, Rachel Dutton

## Abstract

Fermented foods are recognized for their rich microbial diversity and bioactive metabolites, which have recently been linked to anti-inflammatory effects and increased gut microbiome diversity. Despite extensive research on fermented foods including large-scale metagenomic surveys and metabolite characterization, a comprehensive mechanistic understanding of how diverse fermented foods, their microbes, and resulting metabolites interact with human biological pathways remains limited. Here, we systematically profiled over 100 commercially available fermented foods for their potential to prevent inflammation using a human cell-based model. We then generated bulk RNA sequencing of the human cells under these treatment conditions, along with metagenomic sequencing and metabolomics of the fermented foods used in the assays. By generating sample-matched multi-omics data, our work aims to lay the scientific groundwork that will enable scientists to generate predictive models and testable hypotheses about the molecular mechanisms underlying the anti-inflammatory effects of fermented foods. This open-source resource comprising all raw data and parsed intermediary files allows the community to begin elucidating the molecular mechanisms between fermented foods and human biology, paving the way for new, scientifically informed strategies for the user of fermented foods as functional foods.

**Resources:**

- Bioactivity assay results (IFN and NF-kB) of cell-based models of human inflammation with fermented foods applied [available on Zenodo](https://zenodo.org/records/19339682)
- RNA-sequencing of the cell bioactivity assays, including count tables and pathway calculations. Raw sequencing data [available on NCBI](https://www.ncbi.nlm.nih.gov/bioproject/PRJNA1444194), and count tables available on Zenodo
- Shotgun metagenomic sequencing of fermented foods, including species-profiling with a custom fermented food-associated microbial database. Raw sequencing data [available on NCBI](https://www.ncbi.nlm.nih.gov/bioproject/PRJNA1432018), species profiling table [available on Zenodo](https://zenodo.org/records/22086844)
- Untargeted metabolomics of fermented foods catalog through partnership with the Periodic Table of Foods Initiative available on Zenodo

## Background and Summary

Fermented foods are an ancient component of human dietary culture and are increasingly recognized as a rich source of bioactive metabolites that can influence human health. During fermentation, ambient microbial communities or defined starter cultures transform food components into a diverse array of metabolites, including organic acids, alcohols, peptides and other small molecules^1^. Beyond their role in food preservation through the production of organic acids and bacteriocins that inhibit microbial pathogens ^2,3^, fermented food-derived metabolites have been increasingly recognized as potential mediators of human immune and metabolic health^4^. For example, the molecule D-phenyllactic acid (D-PLA), a metabolite produced by lactic acid bacteria and commonly detected in sauerkraut and kimchi, activates human hydroxycarboxylic acid receptor 3 (HCA3) and is hypothesized to play a role in immunomodulation^5–7^. Hundreds of fermented food-associated peptides have been experimentally verified to have anti-hypertensive and anti-oxidative activities^8–10^. Additionally, we recently predicted thousands of fermented-food associated peptide sequences for 17 human health-relevant bioactivity categories using machine learning models^11^.

Despite growing evidence that food-derived metabolites can influence inflammatory responses ^12–14^, characterizing the diversity and abundance of these compounds within a single food remains challenging. Large-scale molecular profiling of foods is often limited by factors such as complex food matrices that hinder extraction, analytical methods that incompletely capture diverse compound classes, and a lack of standardization across laboratories and studies that complicates cross-study comparisons^15,16^. Linking detected molecules to anti-inflammatory activity presents an additional challenge because inflammatory responses are driven by interconnected and context-dependent signaling networks^17^. Although machine-learning approaches are increasingly used to predict anti-inflammatory molecules, their performance is often constrained by limited generalizability due to a lack of broad and diverse training data that are experimentally verified.

Fermented foods have only recently been investigated for their potential anti-inflammatory and immune modulating properties. A human dietary study showed that increased consumption of fermented foods led to decreased circulating markers of inflammation and increased gut microbiome diversity compared to the high fiber-only arm^18^. Anti-inflammatory properties of fermented foods could stem from several sources, including the raw food substrate, activity of specific microbial strains, and metabolites produced during fermentation. Several fermented food-associated microorganisms such as *Lactobacillus plantarum* have been demonstrated to reduce inflammatory markers and alleviate colitis in mice^19–21^. Interestingly, a recent study of metabolite extracts from green cabbage sauerkraut fed to mice were associated with decreased inflammatory markers, increased cecal bacterial CFU counts, and an enrichment of the mucus-associated commensal *Akkermansia muciniphila* compared to the control^22^. This study strongly suggests that fermentation-derived metabolites alone (without the raw food substrate or microorganisms) are sufficient for influencing immune pathways and microbial community dynamics. Despite these observations, a mechanistic understanding of how the different components of fermented foods including food substrates, microbial community composition, and fermentation-derived metabolites contribute to anti-inflammatory responses remains limited.

To address this gap, we profiled over 100 fermented foods for anti-inflammatory activity by applying fermented food extracts to a human cell-based model of inflammation. We generated a paired multi-omics dataset including RNA-sequencing of the treated human cells, and shotgun metagenomic sequencing and metabolomics profiling of the fermented food extracts. The collection comprises more than 100 fermented foods representing 46 different types of fermented foods produced in 21 countries. For each individual food sample, we banked sub-samples for metagenomic shotgun sequencing, untargeted metabolomics data generation, and for preparation of extracts to test for activity in a human cell-based model of inflammation (Figure 1). By linking anti-inflammatory activity in a human cell-based assay with matched metagenomic, metabolomic, and transcriptional profiles, this open resource provides a foundation for identifying candidate bioactive molecules, associated microbial species, and developing predictive models of fermented food-mediated anti-inflammatory activity

**Figure 1.**
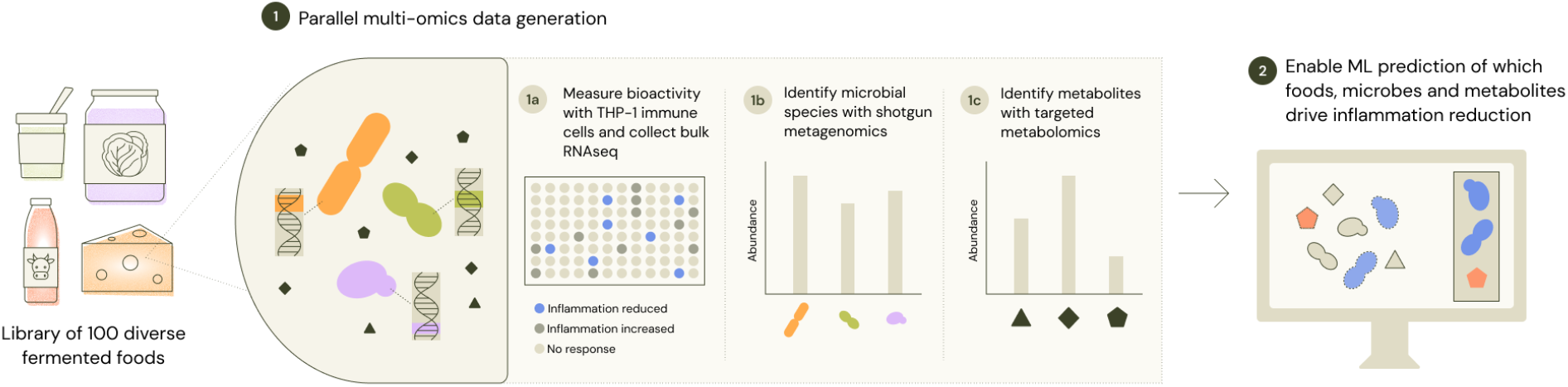
Overview of fermented food sampling and assays. We sampled over 100 fermented foods representing 46 different types of fermented foods produced in 21 countries. Fermented food extracts were applied to human THP-1 dual reporter cells for measuring the activity of IFN and NF-kB reporters. Extract treated cells were also collected for bulk RNA-sequencing. Shotgun metagenomic sequencing and metabolomics were performed on the food extracts. For most samples we were able to collect data for each modality, with all samples resulting in a bioactivity reporter assay result.

### Anti-inflammatory activity of fermented food extracts

We used a human THP-1 dual cell line reporter to measure reporter activity of interferon (IFN) (Figure 2). After filtering normalized IFN activity only for datapoints with cell viability > 70%, most of the fermented food extracts show anti-inflammatory activity (Supplementary Table 1). The top five categories of fermented foods with at least one biological replicate that met this definition were dairy (27 samples), fruit (17 samples), cabbage (14 samples), soy (12 samples), and grain (11 samples).

**Figure 2.**
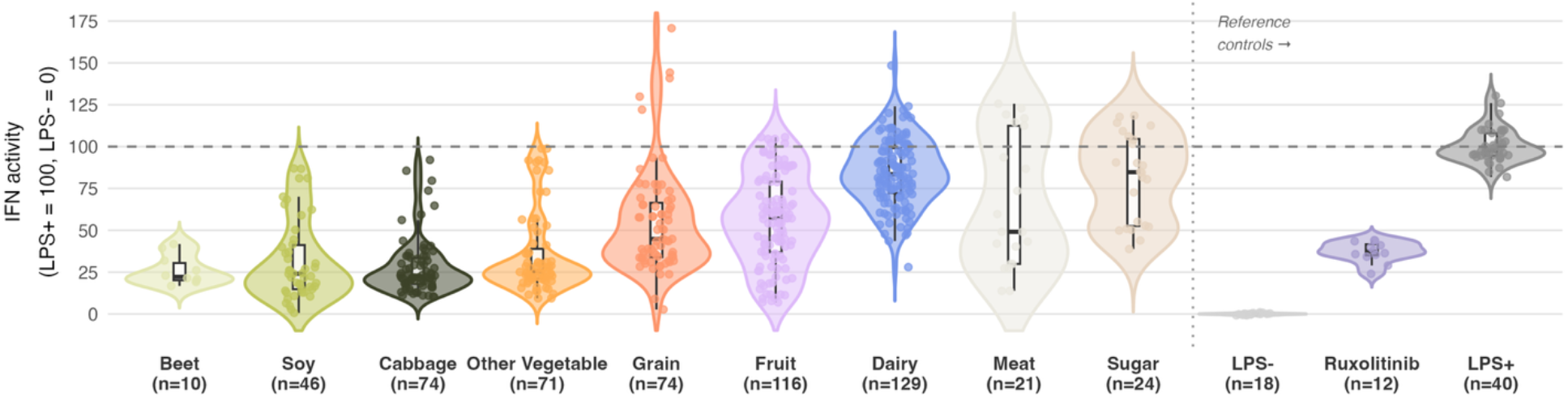
Interferon (IFN) reporter activity of fermented food extracts. Violin plot of IFN activity filtered to only include results with >70% viability across fermented food substrate categories alongside reference controls. Data are normalized relative to the positive control (LPS+ = 100) and negative control (LPS- = 0), indicated by horizontal dashed lines. Individual points represent individual fermented food extracts or control replicates, with sample sizes indicated on the x-axis, and includes biological replicates for a single fermented food extract. Nested boxplots within each violin represent the median and interquartile range (IQR). Reference controls (separated by vertical dotted line) include LPS-(negative control), LPS+ (positive control), and Ruxolitinib (JAK inhibitor, suppressor of IFN signaling).

**Figure 3.**
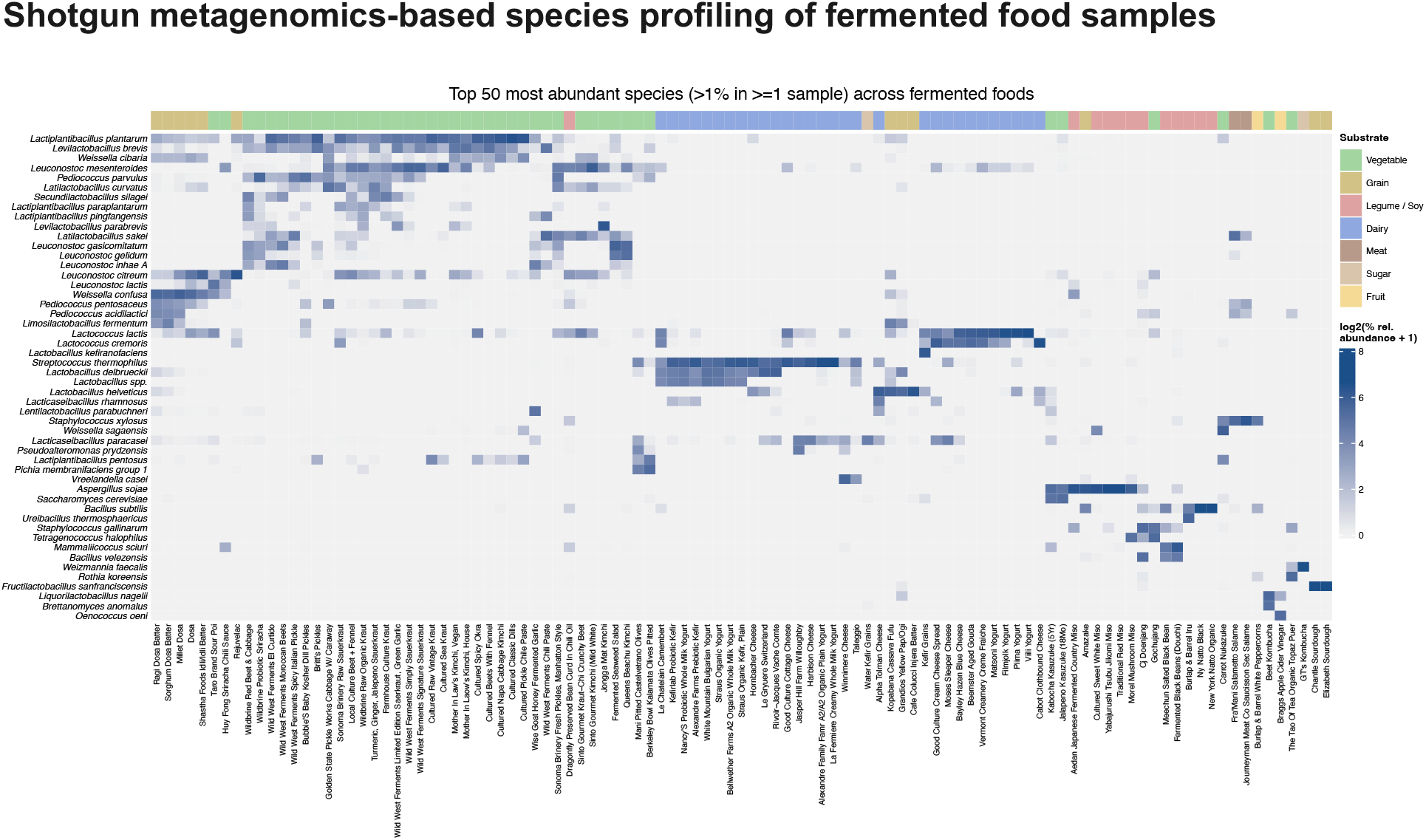
Top 50 most abundant species identified through shotgun metagenomic sequencing. We performed shotgun metagenomic sequencing on most of the fermented foods in our collection that could be successfully extracted. Using a custom fermented food-associated microbial genomes database we previously generated ^11^, we used sylph^25^ to profile each metagenomic sample. The relative abundance of species detected in each metagenomic sample adds up to 100% for all samples, demonstrating the diversity of our custom genome database covers the composition of our samples. The plot above shows the top most 50 abundant species that were present in at least 1 sample and at least 1% relative abundance.

We also measured reporter activity of NF-kB (Supplementary Figure 3) but did not observe detectable anti-inflammatory effects for most of the fermented food extracts, especially compared the positive control PDTC inhibitor of NF-kB. We highly suspect that this dual cell reporter line for NF-kB activity was not sensitive enough for detecting effects of the fermented food extracts. Compared to the recent Caffrey et al. findings that wild sauerkraut lowers NF-kB activity in a GFP THP-1 cell reporter line, the dual reporter cell line may not be appropriate for measuring the activity of fermented food extracts^22^.

We were interested in which samples exhibited the most anti-inflammatory activity by normalized IFN activity, particularly those with similar or lower IFN activity compared to the IFN inhibitor Ruxolitinib. Ruxolitinib is a JAK-inhibitor which block the pro-inflammatory IFN cytokines, and is commonly used for treating myelofibrosis ^23,24^. Samples with low normalized IFN activity that were equal or less than the median normalized IFN reporter activity for Ruxolitinib (normalized median activity of 37.29) were mostly vegetable ferments (20 samples), specifically sauerkraut and pickled vegetables. Other categories included fruit samples (8 samples) such as fermented chili paste, soy samples (8 samples) such as miso, and grain samples (7 samples) such as home-made sourdough and injera.

### Shotgun metagenomics-based species profiling of fermented food samples

We used a custom databased of fermented food-associated microbial genomes that we previously curated from hundreds of publicly available fermented food metagenomes that includes ∼1,300 species-representative genomes comprised of bacterial and eukaryotic species assembled from fermented foods. Shotgun metagenomics was performed on all samples that produced sufficient extracted DNA, and reads were then mapped back onto the custom genome database to determine the presence and abundance of different microbial species within each fermented food. Without filtering the number of hits based on whether the mapped sequencing reads fully covered a genome within the database, we detected 512 distinct species in our dataset. Every sample totaled 100% sequence coverage from the detected hits, demonstrating that our custom genome database was sufficiently diverse to cover the sequence diversity within our samples. When filtering the hits for genomic sequence coverage of at least 1X, this greatly reduces the total number of species detected to 226. In particular, this affects detected distinct eukaryotic species from 36 species to 12, suggesting that increased sequencing depth may be needed to cover the larger eukaryotic genomes. With the filter applied, the top three most detected bacterial species are *Lactobacillus plantarum* (40 samples), *Lactococcus lactis* (37 samples), and *Leuconostoc mesenteroides* (34 samples). With the filtered applied, only the eukaryotic species *Aspergillus sojae* and *Saccharomyces cerevisiae* are detected in more than one sample.

### Untargeted metabolomic profiling of fermented food samples

A highly differentiated aspect of our dataset is the high-quality metabolomics results we achieved through the Periodic Table of Foods Initiative (PTFI) standardized platform.Untargeted LC-MS/MS metabolomics generates large volumes of detected signal that cannot be annotated using existing public spectral libraries, leaving most features in complex biological matrices uncharacterized. To address this, the PTFI and Verso Bioscience developed MarkerLab, a standardized platform that aligns the retention time dimension of LC-MS/MS instruments across a large multi-laboratory network and consolidates MS1 isotope and adduct envelopes to assign a molecular formula to each detected feature. The primary output of the MarkerLab pipeline is a formula–retention index (RI) tuple for each compound with attached MS/MS spectra and sample provenance metadata. The addition of cross-laboratory retention index reproducibility and formula-level constraint substantially improves annotation accuracy relative to spectral matching alone, reducing false annotations arising from isobaric compounds and spectral ambiguity. Using this platform, the PTFI and Verso have developed a reference library of compounds observed reproducibly across the network, currently comprising more than 3,000 structurally identified compounds and more than 25,000 unknown compounds, each assigned to a stable identifier. This library is used to annotate new samples analyzed by the network while simultaneously being expanded by new observations and new information about the structural identity of the unknown compounds.

We profiled metabolites in our fermented food collection using reverse phase (Figure 4) and hydrophilic interaction liquid chromatography (HILIC) (Figure 5) methods. In the reverse phase data, we were able to detect 454 distinct metabolites, and in the HILIC data we were able to detect 7799 distinct metabolites with 4347 of those metabolites detected with high confidence and 1113 named compounds with high confidence. Another strength of using the standardized pipeline through the PTFI is the ability to compare unnamed compounds (given PTF or NP IDs) that may or may not have an identified structure associated with it across different food samples that the PTFI has analyzed. This resource represents a high-quality, standardized platform for analyzing food metabolites, which have previously only had few large-scale efforts such as the Global Food Omics study ^26^.

**Figure 4.**
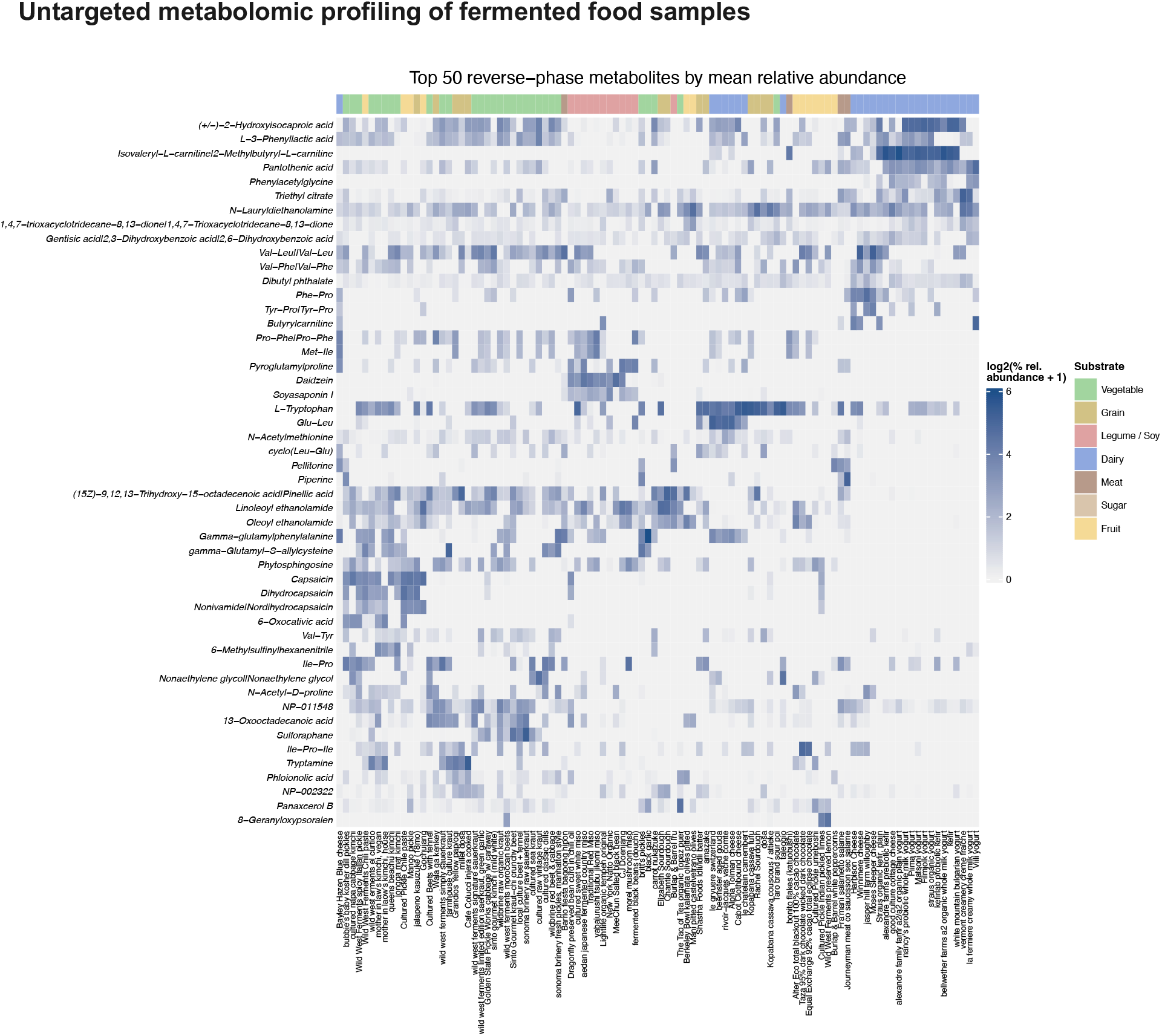
Top 50 most abundant metabolites by mean relative abundance detected by reverse-phase metabolomics. Each row represents the log2 normalized relative abundance of a metabolite in different samples. Due to the detection and normalization limits of the metabolomics method, abundance of metabolites should only be compared in different samples for a single metabolite, and not across different metabolites.

**Figure 5.**
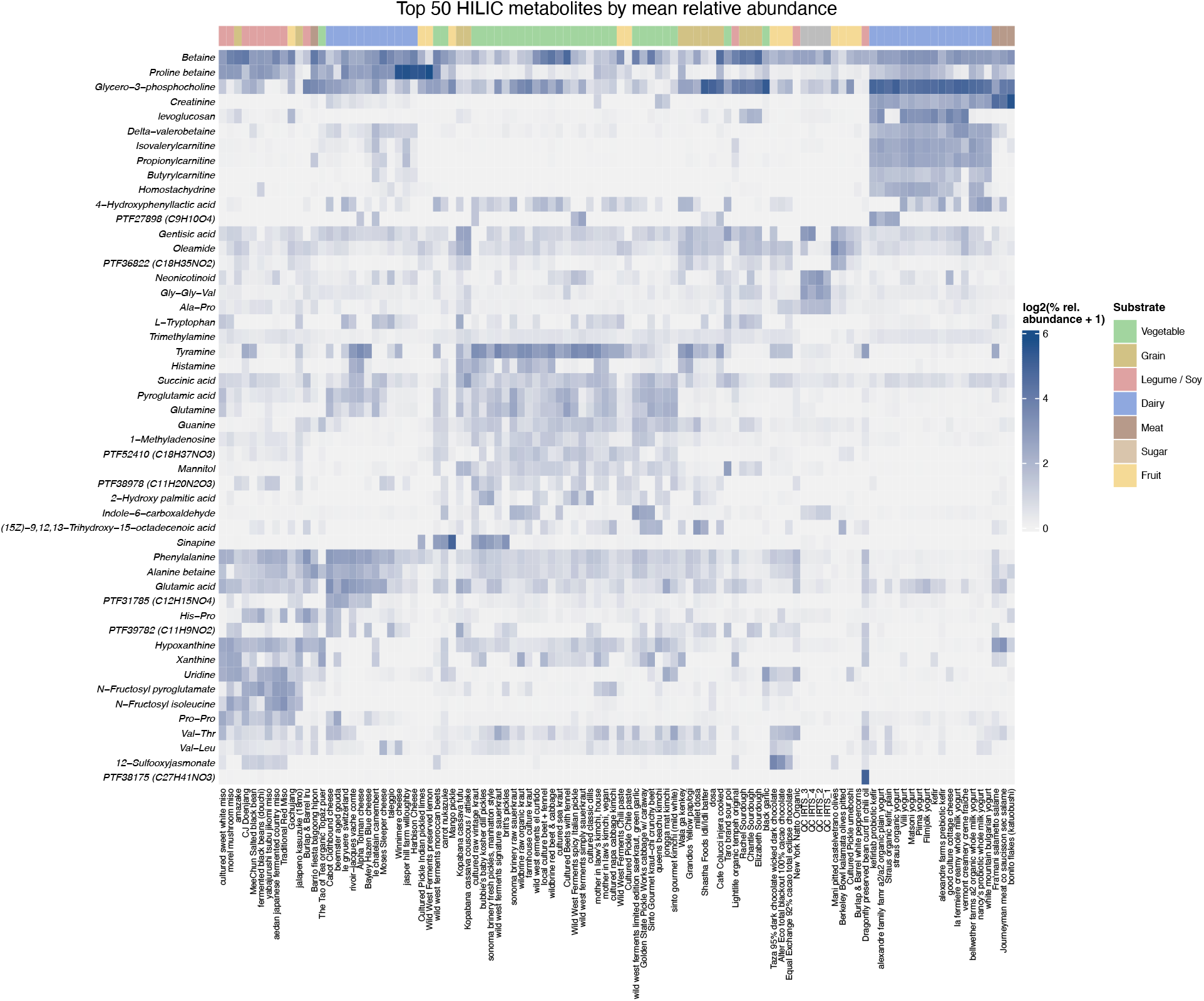
Top 50 most abundant metabolites by mean relative abundance detected by HILIC metabolomics. Each row represents the log2 normalized relative abundance of a metabolite in different samples. Due to the detection and normalization limits of the metabolomics method, abundance of metabolites should only be compared in different samples for a single metabolite, and not across different metabolites. Compounds that start with PTF are unnamed compounds that have been identified in other PTFI datasets.

Specifically for fermented foods, we were able to identify several metabolites that have been shown to be associated with human health-relevant pathways. Particularly, aryl-lactates such as D-phenyllactic acid, indole-3-lactic acid, and 4-hydrophenyllactic acid have been shown to be present across diverse commercially available fermented foods and suggested to interact with immunomodulatory receptors including the human hydrocarboxylic acid receptor (HCAR3) and the aryl hydrocarbon receptor (AhR) ^27–29^. D-phenyllactic acid (D-PLA), which has previously been detected in high concentrations in sauerkraut and kimchi^5^, was detected in 92% of the samples in our dataset. We also detected 4-hydroxyphenyllactic acid in 53% of samples, most commonly in vegetable ferments (20 samples) including pickled vegetables, sauerkraut, kimchi, and dairy (18 samples) including kefir, yogurt, and cheese. We detected indole-3-lactic acid in 22% of samples, mostly in vegetable ferments (8 samples) such as kimchi and sauerkraut, and dairy (9 samples) including cheese, yogurt, and kefir.

### Summary

Overall, this dataset provides an integrated resource for connecting the microbial and chemical diversity of fermented foods to effects on human inflammatory pathways. By pairing cell-based bioactivity measurements with matched transcriptional responses, metagenomic profiles, and standardized metabolomics across a broad collection of commercially available fermented foods, this work enables the field to start building predictive models of fermented foods and their effects on human health.

## Methods

### Food sampling

Fermented foods were collected from local supermarkets, online suppliers, and home-made products. A complete list of the sampled fermented foods and their associated metadata including sampling date and location can be found at https://zenodo.org/records/22086559 under the FF samples – Combined master sheet.tsv. Once each food sample was acquired, the sample was split between 50 mL conical tubes stored at 4°C for future sampling and reference, cryovial tubes stored at -80°C for use in the bioassay, cryovial tubes with Zymo DNA/RNA shield and shipped for metagenomic sequencing or shipped on dry ice for metabolomics measurements through the Periodic Table of Foods Initiative.

### Shotgun metagenomic sequencing and analysis

Shotgun metagenomic sequencing was performed at SeqCoast Genomics. Briefly, samples were transferred to MagMAX Microbiome Bead Beating Tubes (FisherSci #A42351) and then mixed with Qiagens CD1 Lysis Buffer (from the Qiagen DNeasy 96 PowerSoil Pro QIAcube HT Kit [#47021]). Bead beating was then performed using a Vortex Genie 2 for 40 minutes. After bead-beating lysis, samples were extracted using the Qiagen DNeasy 96 PowerSoil Pro QIAcube HT Kit (#47021).

DNA samples were prepared for whole-genome sequencing using the Illumina DNA Prep tagmentation kit (#20060059) with Illumina Unique Dual Indexes. Sequencing was performed on the Illumina NextSeq2000 platform using a 300-cycle XLEAP-SBS flow cell kit to produce 2x150bp paired reads. 1-2% PhiX control was spiked into the run to support optimal base calling. Read demultiplexing, read trimming, and run analytics were performed using DRAGEN v4.2.7, an on-board analysis software on the NextSeq2000.

Raw reads were processed and mapped to a custom fermented food-associated microbial genome database using the metagenome-containment-profiler Nextflow workflow (available on GitHub at https://github.com/MicrocosmFoods/metagenome-containment-profiler). Briefly, a custom microbial genome database was constructed from publicly available fermented food genome databases and de-replicated at 95% ANI (McDaniel et al. 2026). The raw genome database is available on Zenodo at https://zenodo.org/records/15794524. The pipeline filters raw FASTQ reads and assigns reads to the input database using sylph, a k-mer containment-based algorithm for read assignment ^25^.

### Metabolomics

Whole food samples were processed through the Periodic Table of Food Initiatives (PTFI) protocols for reverse phase LC-MS/MS metabolomics and polar metabolomics (HILIC LC-MS/MS) at a PTFI Center for Excellence site. Polar and semi-polar metabolites were analyzed using the MarkerLab HILIC LC-MS/MS platform. Samples are prepared starting with 30 µl of serum and uses an MTBE-based biphasic extraction to obtain a clean aqueous (polar) fraction, largely free of phospholipids. The extraction protocol is modified based on Matyash et al. ^30^, and the LC-MS method is based on Folz et al ^31^. The reverse phase metabolomics method relies on a standard LC-MS protocol referenced in Odenkirk et al ^15^.

### Cell-based assays of human inflammation in human THP-1 cells

Human THP1-Dual cells (InvivoGen) were maintained in RPMI 1640 supplemented with GlutaMAX, 25mL HEPES, 10% (v/v) fetal bovine serum (FBS), and 1% penicillin-streptomycin at 37C with 5% CO_2_. Viable cells were quantified using a Countess II FL automated cell counter, harvested, and centrifuged at 300 x g for 7 minutes. The supernatant was aspirated, and the cell pellet was resuspended in pre-warmed RPMI 1640 + GlutaMAX containing 10% (assay medium) to a final concentration of 1E6 cells/mL. Aliquots of the cell suspension 100µL/well were seeded into clear, flat-bottom 96-well assay plates. Reconstituted food extracts and experimental controls 50µL were added to designated wells, and plates were incubated at 37C with 5% CO_2_ for 2 hours. Following pre-treatment, cells were stimulated by adding 50µL of 4x lipopolysaccharide (LPS) prepared in assay medium to achieve an in-well concentration of 250ng/mL. Negative control wells received 50µL of assay medium alone. Plates were subsequently incubated at 37C with 5% CO_2_ for 24 hours.

Following incubation, plates were centrifuged at 300 x g, and 101µL of supernatant was harvested for downstream analysis. Cell viability was determined directly in the treatment plates by adding 1µL of a combined 1-methoxy-phenazine methosulfate (1-methoxy-PMS) and WST-8 solution to yield final in-well concentrations of 200µM and 500µM, respectively. Plates were incubated at 37C with 5% CO_2_ for 2 hours, and absorbance was measured at 450nm using a SpectraMax 340PC microplate reader.

NF-kB inducible secreted embryonic alkaline phosphatase (SEAP) activity was measured using QUANTI-Blue Solution (InvivoGen) prepared by mixing 100x QB reagent and 100x QB buffer in sterile water. Working solution 90µL was added to a clear flat-bottom assay plate, followed by 10µL of harvested culture supernatant. After incubation at 37C with 5% CO_2_ for 30 minutes, absorbance was recorded from 620nm to 655nm at 5nm increments on a SpectraMax 340PC microplate reader.

IFN-inducible reporter luciferase activity was measured using QUANTI-Luc 4 (InvivoGen). Working solution 50µL, prepared by combining 20x QUANTI-Luc 4 Reagent and 25x QUANTI-Luc 4 Stabilizer in sterile water, was added to opaque white flat-bottom plates, followed by harvested supernatant. Luminescence was measured immediately using a Tecan Infinite Evolution microplate reader.

### RNA-sequencing and analysis of human THP-1 cells treated with fermented food extract

Human THP1-Dual cells were treated with fermented food extracts and LPS or controls following the workflow described above. Treated cells were collected and preserved in Zymo DNA/RNA Shield according to manufacturer specifications for commercial sequencing (Plasmidsaurus). RNA extraction, library prep, and sequencing were performed at Plasmidsaurus. Briefly, total RNA was isolated via a bead-based extraction method. RNA concentrations were quantified using a fluorescence-based microplate assay and normalized prior to library construction. Poly(A) mRNA was converted to cDNA via reverse transcription and second-strand synthesis, followed by tagmentation, library indexing, and PCR amplification. Sequencing was performed to a target depth of 1E7 reads per sample with an average read length of 90bp.

Reads were quality filtered using fastp v0.24.0^32^ with poly-X tail trimming, 3 quality-based tail trimming, a minimum Phred quality score of 15, and a minimum length requirement of 50 bp. Read quality before and after filtering was assessed with FastQC v0.12.1^33^. Quality-filtered reads were aligned to the human reference genome assembly GRCh38 at assembly accession GCA_000001405.29 using STAR aligner v2.7.11^34^ with non-canonical splice junction removal and output of unmapped reads, followed by coordinate sorting with samtools v1.22.1. PCR and optical duplicates were removed using UMI-based deduplication with UMIcollapse v1.1.0^35^.

Gene-level expression quantification was performed using featureCounts (subread package v2.1.11)^36^ with strand-specific counting, multi-mapping read fractional assignment, exons and three prime UTR as the feature identifiers, and grouped by gene_id. Final gene counts were annotated with gene biotype and other metadata extracted from the reference GTF file. Further technical documentation for library prep and initial bioinformatics is described on the Plasmidsaurus website. (https://plasmidsaurus.com/technical-documentation/rna.)

## Data availability

Raw metagenomic sequencing files are available on the NCBI SRA at BioProject accession PRJNA1432018. Raw RNA sequencing files are available on the NCBI SRA at BioProject accession PRJNA1444194. Raw and normalized bioactivity measurements of IFN and SEAP from THP-1 cell assays are available on Zenodo at https://zenodo.org/records/18969859. Raw counts and counts per million for the bulk RNAseq experiments are available on Zenodo at https://zenodo.org/records/20618557. Result tables for the reverse phase and HILIC metabolomics methods are available on Zenodo https://zenodo.org/records/22086559. All code and workflows are openly available under an MIT license on GitHub at https://github.com/MicrocosmFoods, and the supermarket sweep GitHub repository https://github.com/MicrocosmFoods/supermarket-sweep.

## Acknowledgements

We would like to thank Jordan Toutounchian for consulting on bioassay methods, and Jas Neal and Micaela Caffrey for the overview figure visualization work. Metabolomics was performed by the Periodic Table of Foods Initiative (PTFI) through a coordinated network of labs, including Verso Bio. We would like to thank Steven Watkins at Verso Bio for providing assistance in metabolomics data interpretation and writing. This research was funded by the Astera Institute as part of Rachel Duttons residency project. Additional support for metabolomics was received from the Periodic Table of Foods Initiative.

## Supplementary Material

**Supplementary Figure 1.**
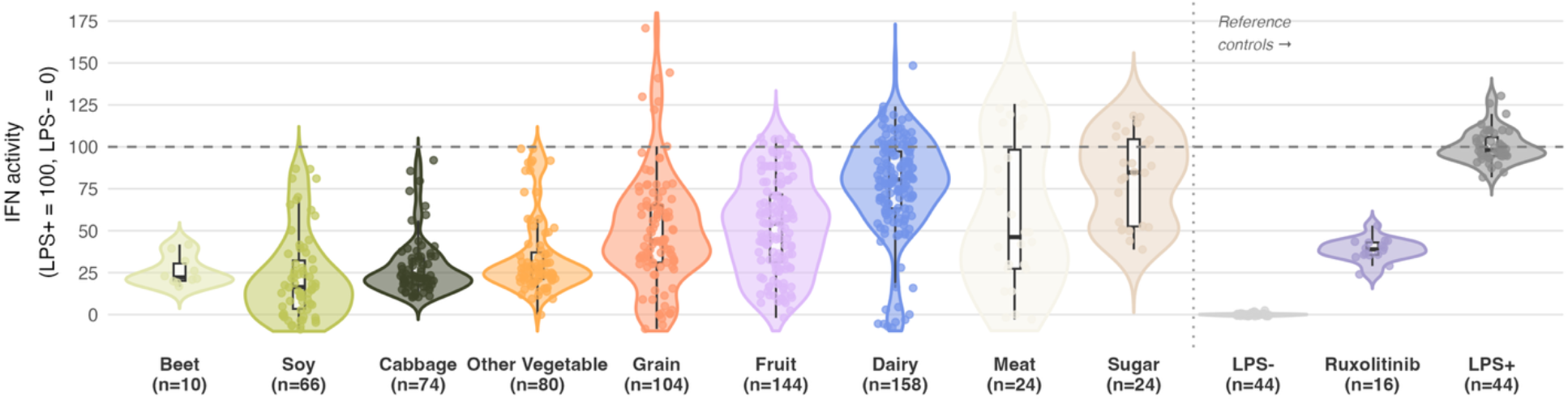
Unfiltered Interferon (IFN) reporter activity. Violin plots depicting normalized IFN activity across fermented food substrate categories alongside reference controls. Data are normalized relative to the positive control (LPS+ = 100) and negative control (LPS- = 0), indicated by horizontal dashed lines. Individual points represent individual fermented food extracts or control replicates, with sample sizes indicated on the x-axis. Nested boxplots within each violin represent the median and interquartile range (IQR). Reference controls (separated by vertical dotted line) include LPS-(negative control), LPS+ (positive control), and Ruxolitinib (JAK inhibitor, suppressor of IFN signaling).

**Supplementary Figure 2.**
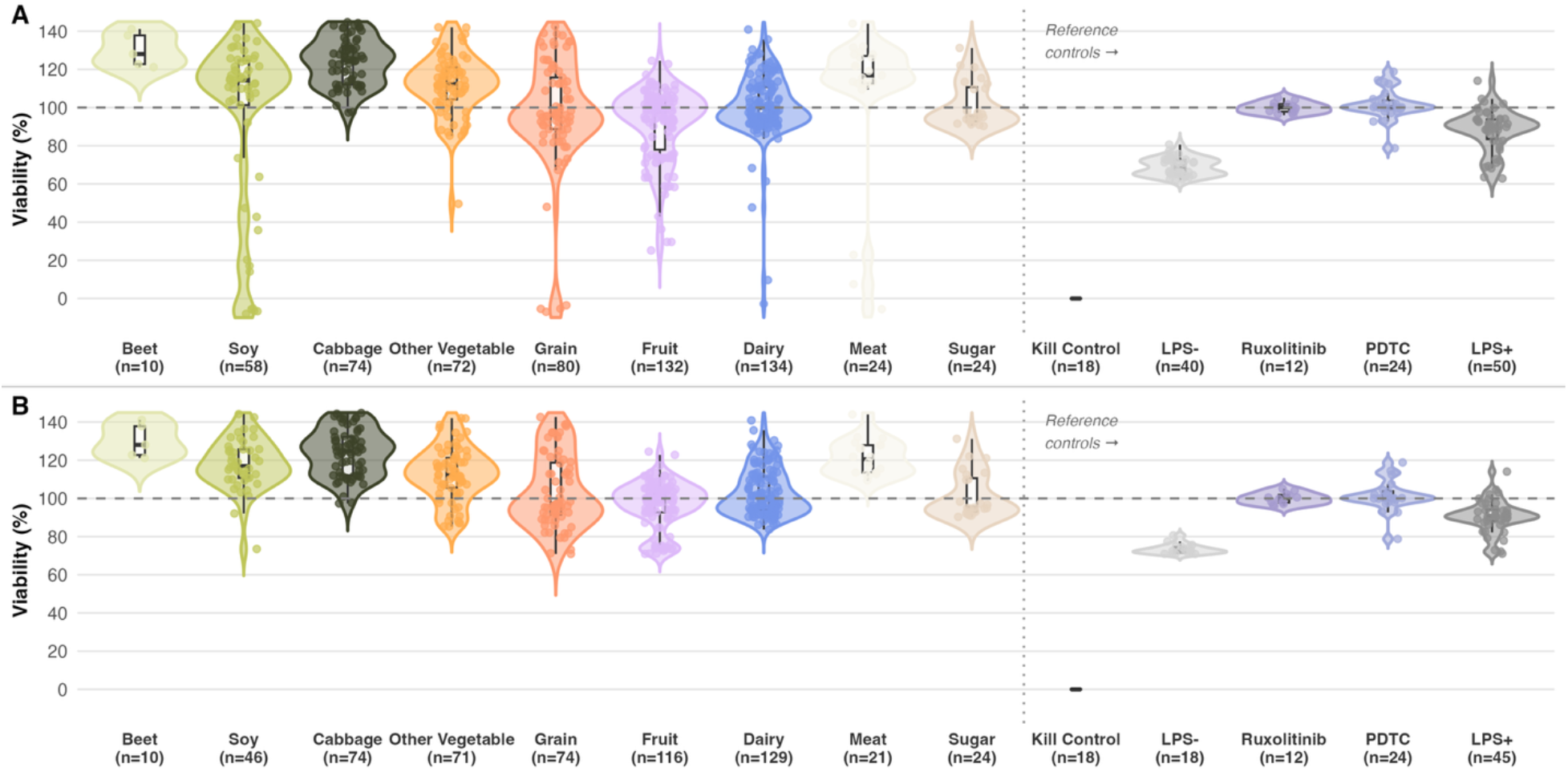
WST-8 Viability measurements. Cellular viability and metabolic activity were measured in THP-1 cells treated with fermented food extracts or controls using the WST-8 assay. This assay quantifies intracellular dehydrogenase and mitochondrial enzyme activity, which correlates directly with cell viability. Viability values (%) were normalized relative to baseline, where 100% represents the untreated vehicle control baseline (indicated by the horizontal dashed line) and 0% represents the Kill Control condition (cells treated with a cytotoxic agent to induce complete cell death). Individual points represent distinct fermented food samples or control replicates (indicated on the x-axis), and nested boxplots represent the median and interquartile range (IQR). Values exceeding 100% (e.g., in Beet, Cabbage, Dairy, and Meat) reflect potential metabolic hyperactivation or cellular proliferation, which might be fueled by nutrient-rich fermentation components such as fermentable sugars, organic acids, or amino acids that directly feed mitochondrial dehydrogenases. Conversely, values below baseline indicate metabolic suppression or cytotoxicity, potentially due to high acidity and pH shifts, high osmolality and salinity, or cytotoxic byproducts. To ensure that downstream pathway readouts were not confounded by compromised cell health, samples exhibiting viability below 70% were excluded from downstream analyses. Reference controls (separated by the vertical dotted line) include Kill Control (complete cell death baseline, ∼0% viability), LPS-(negative control), Ruxolitinib (JAK inhibitor), PDTC (NF-kB inhibitor), and LPS+ (positive control).

**Supplementary Figure 3.**
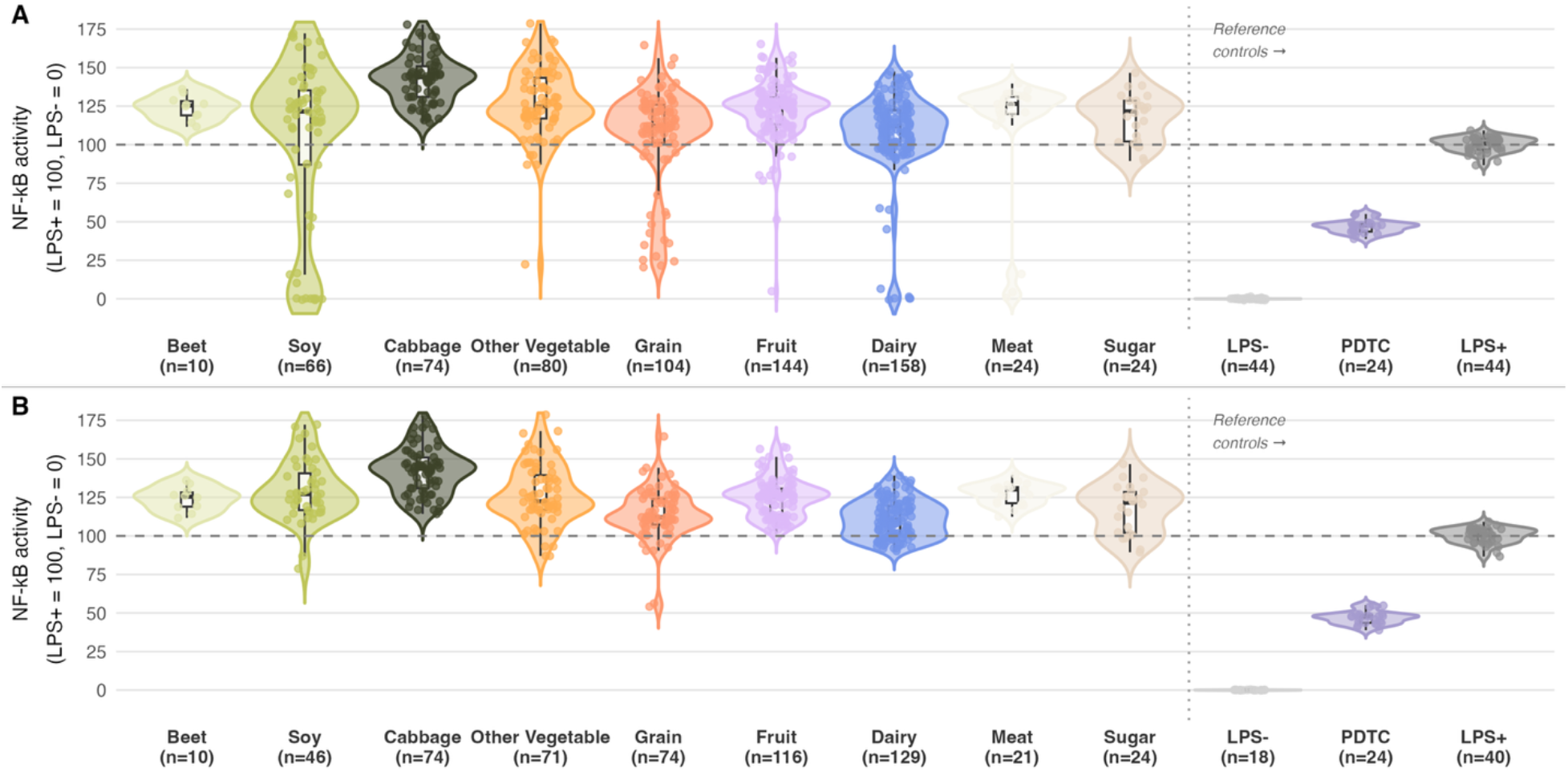
Unfiltered and filtered NF-kB reporter activity. NF-kB reporter activity unfiltered (A) and filtered for results with normalized viability > 70% (B). Results are normalized relative to the positive control (LPS+ = 100) and negative control (LPS- = 0), indicated by horizontal dashed lines. Individual points represent individual fermented food extracts or control replicates, with sample sizes indicated on the x-axis. Nested boxplots within each violin represent the median and interquartile range (IQR). Reference controls (separated by vertical dotted line) include LPS-(negative control), LPS+ (positive control), and PDTC (ammonium pyrrolidinedithiocarbamate, an inhibitor of NF-kB signaling).

**Supplementary Table 1.**
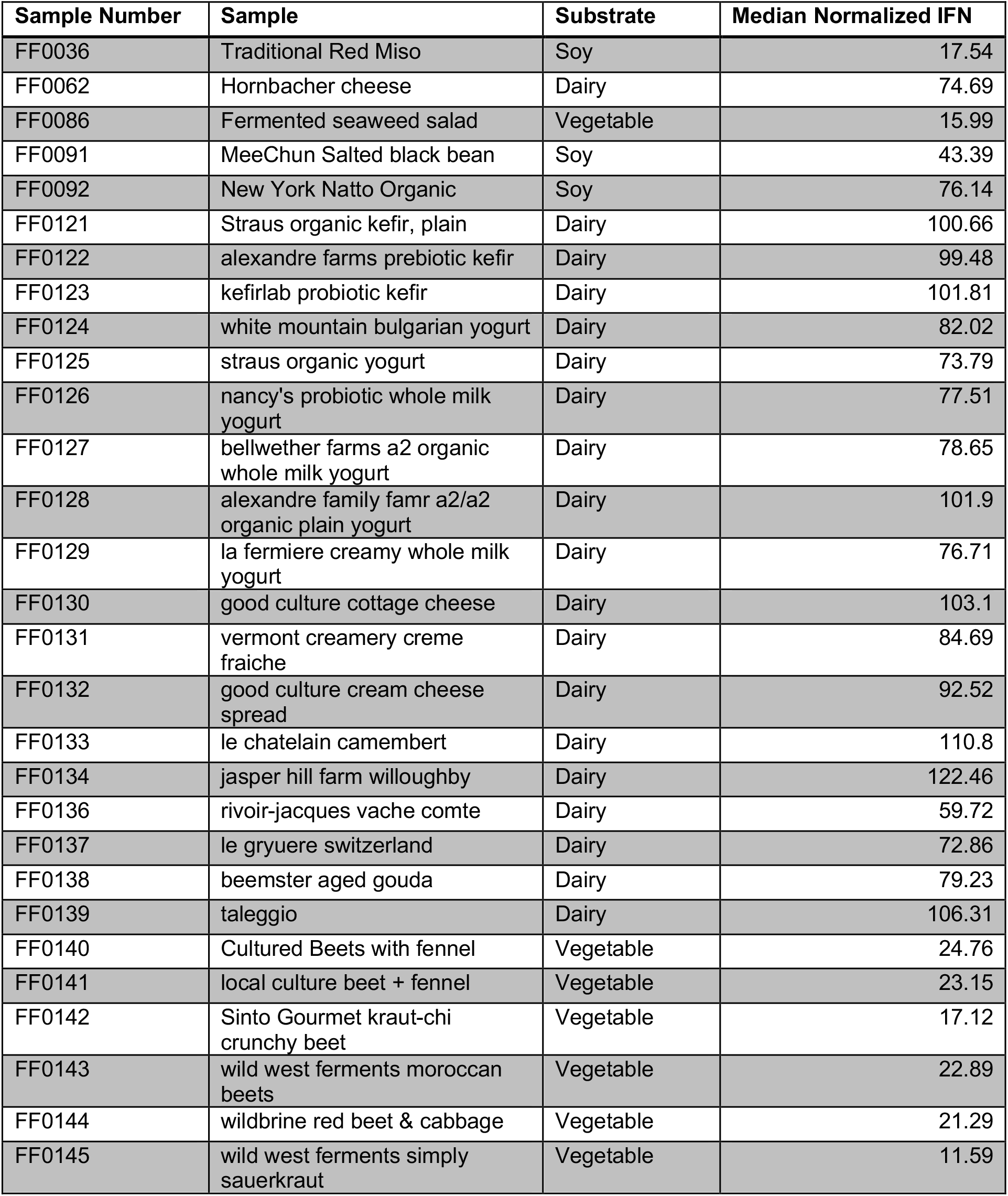
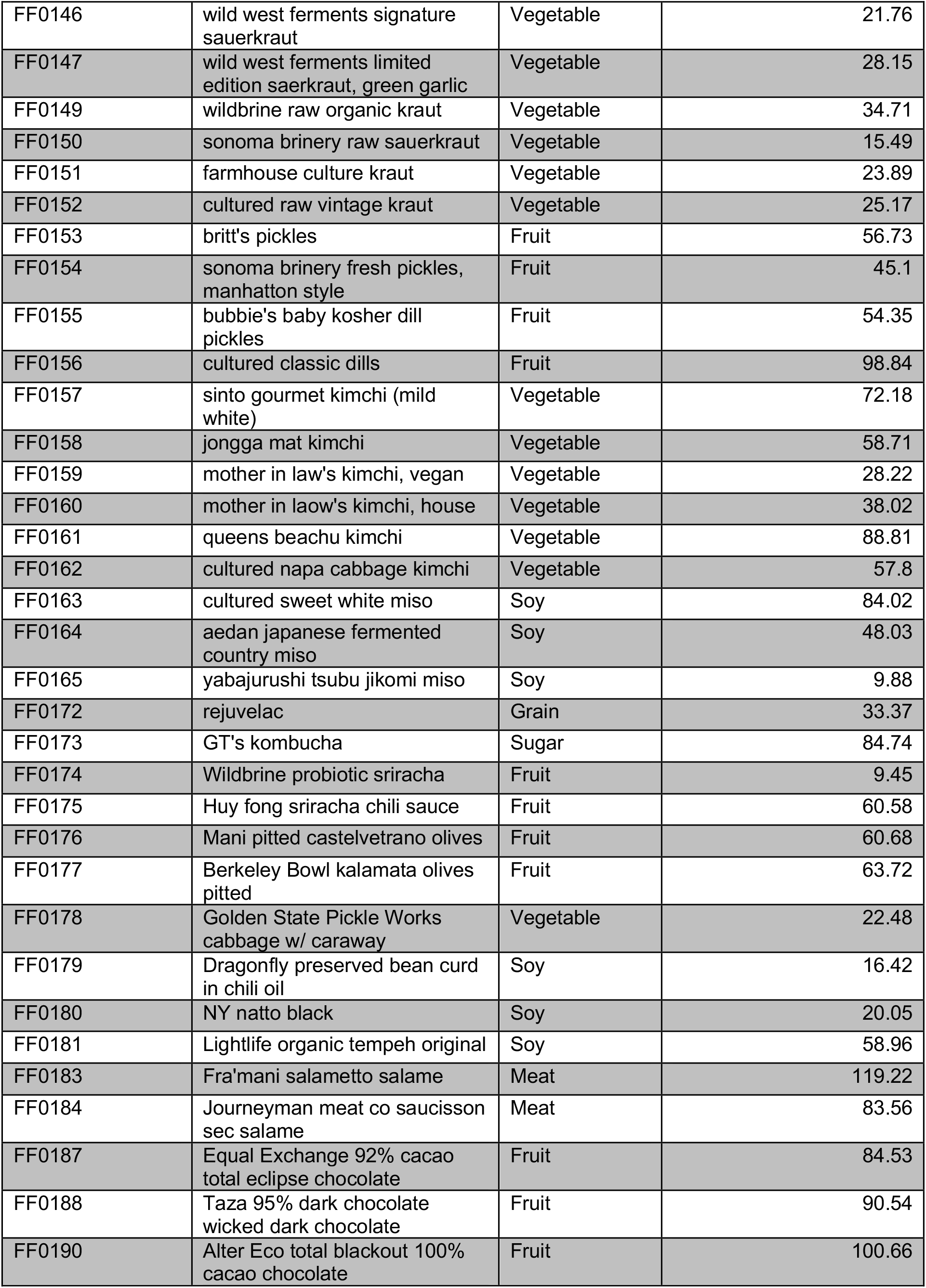
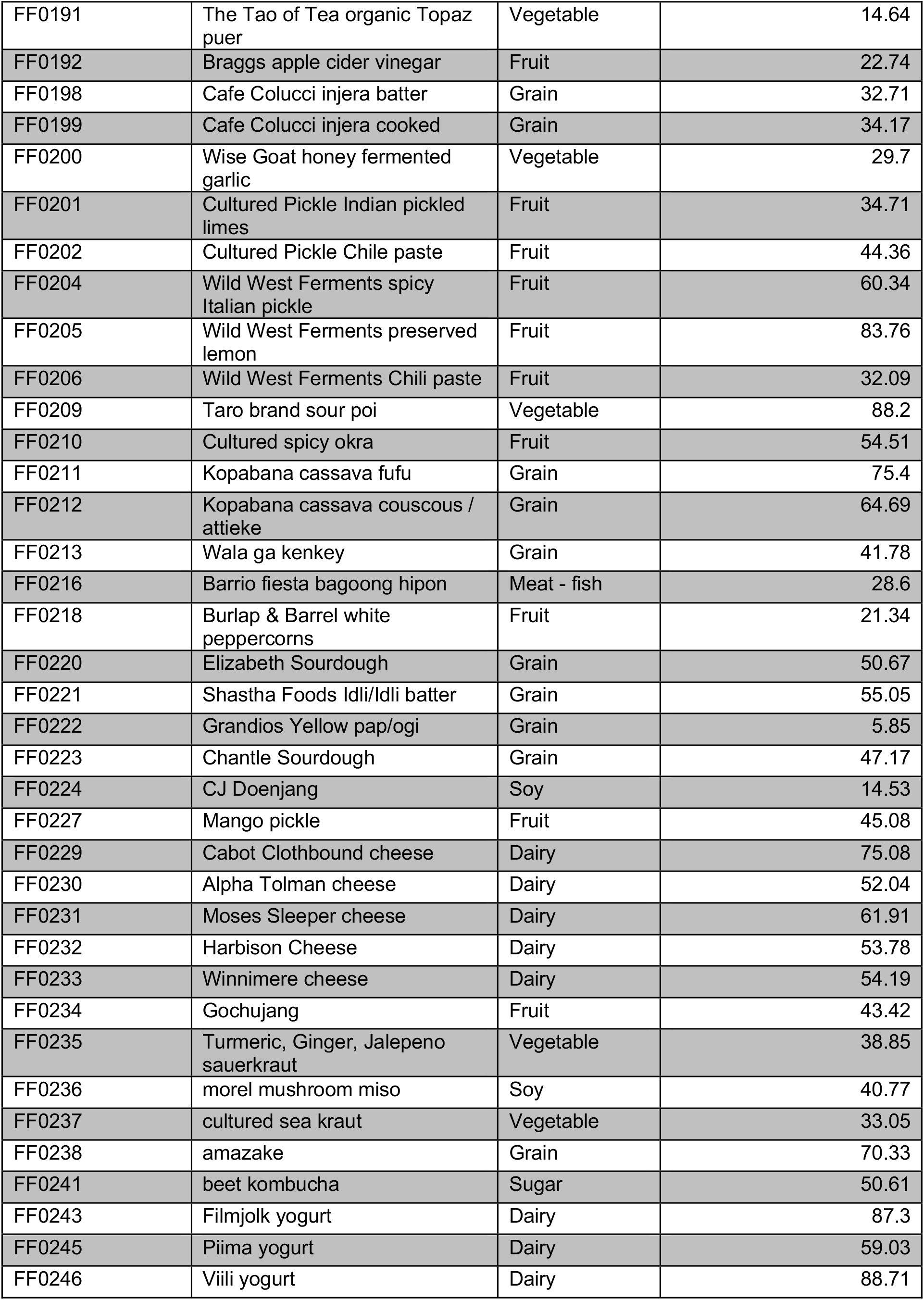
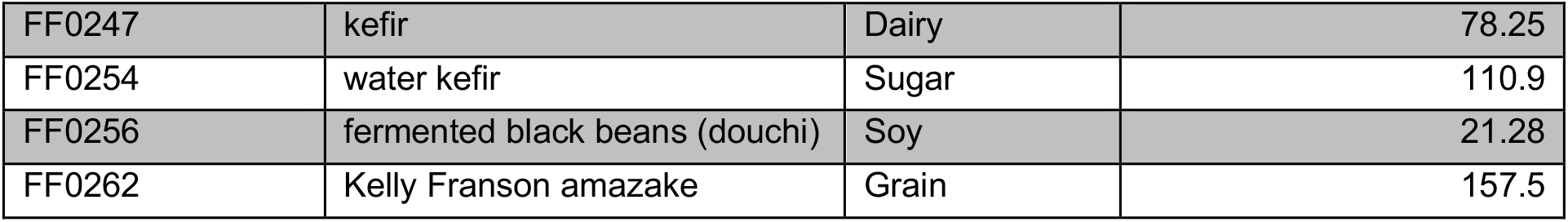
Median normalized IFN for each fermented food sample, where results were included for data points where cell viability > 70%.

